# RNA interactome of hypervirulent *Klebsiella pneumoniae* reveals a small RNA inhibitor of capsular mucoviscosity and virulence

**DOI:** 10.1101/2024.06.23.600155

**Authors:** Kejing Wu, Xingyu Lin, Yujie Lu, Rui Dong, Hongnian Jiang, Sarah L Svensson, Jiajia Zheng, Ning Shen, Andrew Camilli, Yanjie Chao

**Affiliations:** Microbial RNA Systems Biology Unit, Center for Microbes, Development and Health (CMDH), Shanghai Institute of Immunity and Infection, Chinese Academy of Sciences, Shanghai, 200031, China; University of Chinese Academy of Sciences, Beijing, 100049, China; Key Laboratory of RNA Innovation, Science and Engineering (RISE), Shanghai Institute of Biochemistry and Cell Biology, Center for Excellence in Molecular Cell Science, Chinese Academy of Sciences, Shanghai, 200031, China; Department of Laboratory Medicine and Department of Pulmonary and Critical Care Medicine, Center of Infectious Disease, Peking University Third Hospital, Beijing, 100191, China; Department of Molecular Biology and Microbiology, Tufts University School of Medicine, Boston, Massachusetts 02111. USA

**Keywords:** *Klebsiella pneumoniae*, CR-HvKP, sRNA, CRP, MlaA, Fbp, capsule, hypermucoviscosity, iRIL-seq

## Abstract

Hypervirulent *Klebsiella pneumoniae* (HvKP) is an emerging bacterial pathogen causing invasive infection in immune-competent humans. The hypervirulence is strongly linked to the overproduction of hypermucovisous capsule, but the underlining regulatory mechanism of hypermucoviscosity (HMV) has been elusive, especially at the post-transcriptional level mediated by small noncoding RNAs (sRNAs). Using a recently developed RNA interactome profiling approach, we interrogate the Hfq-associated sRNA regulatory network and establish the intracellular RNA-RNA interactome in HvKP. Our data reveal numerous interactions between sRNAs and HMV-related mRNAs, and identify a plethora of sRNAs that repress or promote HMV. One of the strongest repressors of HMV is ArcZ, which is activated by catabolite regulator CRP and targets many HMV-related genes including *mlaA* and *fbp*. We discover that MlaA and its function in phospholipid transport is crucial for capsule retention and HMV, inactivation of which abolished *Klebsiella* virulence in mice. ArcZ overexpression significantly reduced bacterial burden in mice and reduced HMV in multiple carbapenem-resistant and hypervirulent clinical isolates with diverse genetic background, indicating it is a potent RNA inhibitor of bacterial pneumonia with therapeutic potential. In summary, our work unravels a comprehensive map of the RNA-RNA interaction network of HvKP and identifies a novel CRP-ArcZ-MlaA regulatory circuit of HMV, providing mechanistic insights into the posttranscriptional virulence control in a superbug of global concern.

**HIGHLIGHTS:**

- Global RNA-RNA interactome map in hypervirulent *Klebsiella pneumoniae*
- Hfq and multiple small RNAs regulate capsular hypermucoviscosity
- ArcZ targets *mlaA* that is required for lipid transport, hypermucoviscosity and virulence
- Crp is a transcriptional activator of ArcZ governing a new virulence circuit

## INTRODUCTION

The Gram-negative *Klebsiella pneumoniae* is an emerging bacterial pathogen in the era of global antibiotic crisis (Wyres et al., 2020). As a member of the ESKAPE pathogens, antibiotic-resistant *K. pneumoniae* is a leading cause of nosocomial infections and is difficult to treat by frontline drugs including carbapenems (Z. Liu et al., 2023; Pendleton et al., 2013; Rice, 2010). Whereas most classical *K. pneumoniae* strains affect immuno-deficient populations, a distinct pathotype of hypervirulent *K. pneumoniae* (HvKP) emerged in Asia in the 1980s that mainly infects young and healthy people, causing community-acquired pneumonia, liver abscess, endophthalmitis and meningitis (Liu et al., 1986; Piednoir et al., 2020; Zou and Li, 2021). More concerningly, HvKP can evolve to carbapenem-resistant (CR) strains via mobilizing plasmids, giving birth to CR-HvKP superbugs that are responsible for bursting endemics in China that pose urgent threats to global public health and biosecurity (Tian et al., 2022; Yang et al., 2022; Zhang et al., 2016; Zhu et al., 2022).

The virulence of HvKP is strongly linked to overproduction of capsular polysaccharides anchored to outer membrane (Walker and Miller, 2020). With a limited number of recognized virulence traits such as lipopolysaccharide, fimbriae and siderophores, HvKP has been best characterized by its hypermucoviscosity (HMV) phenotype (Paczosa and Mecsas, 2016). HvKP can form elastic strings longer than 5 mm when lifting their colonies from agar plates, a visible feature used in clinical diagnosis (Zhu et al., 2022). HMV is largely attributed to the over-production of capsular and is required for invasive infections and pathogenesis (Walker and Miller, 2020). The emergence of HMV has been associated with lateral acquisition of new regulators of capsule polysaccharides (Cheng et al., 2010; Lai et al., 2003), such as the *rmpADC* genes that promote capsule biosynthesis, transmembrane export and chain extension (Ovchinnikova et al., 2023; Walker et al., 2020, 2019).

Besides the capsule biosynthesis gene cluster, dozens of other genes have been found by global genetic screens to promote or suppress HMV (HMV-related genes), belonging to various pathways including central metabolism, iron homeostasis, RNA degradation, lipid transport, as well as global transcription factors such as CRP and OmpR (Dorman et al., 2018; Lin et al., 2013; Mike et al., 2021; Ou et al., 2017; Wang et al., 2023). Mounting evidence supports that HMV is tightly controlled at multiple layers by an as yet undefined regulatory network in *K. pneumoniae*.

Post-transcriptional control of HMV at the RNA level has been implicated in *K. pneumoniae*, but with little mechanistic understanding. For example, the major RNA chaperone Hfq has been shown to regulate capsule production and pathogenesis (Chiang et al., 2011; Huang et al., 2012). Hfq, as a conserved RNA-binding protein found in most bacteria, functions to stabilize small regulatory RNAs (sRNAs) and facilitate their base-pairing with target mRNAs (**Figure 1A**) (Holmqvist and Vogel, 2018; Hör et al., 2020; Vogel and Luisi, 2011). Hfq and its associated sRNAs regulate hundreds of mRNAs up to a quarter of the transcriptome in models such as *E. coli* and *Salmonella*, forming large post-transcriptional regulatory networks mediated by direct RNA-RNA interactions (Liu et al., 2023; Papenfort and Melamed, 2023; Svensson and Chao, 2022). These RNA regulatory networks have been shown to control virulence programs, metabolic fluxes and stress responses in a wide range of bacterial pathogens including *S. enterica*, *Pseudomonas aeruginosa, Vibrio cholerae,* and *Yersinia pestis* (Caldelari et al., 2013; Chao and Vogel, 2010; Hör et al., 2020; Papenfort and Melamed, 2023; Wang et al., 2020). However, the nature of this Hfq-mediated RNA network in the emerging hypervirulent pathotype of *K. pneumoniae* is little understood.

**Figure 1.**
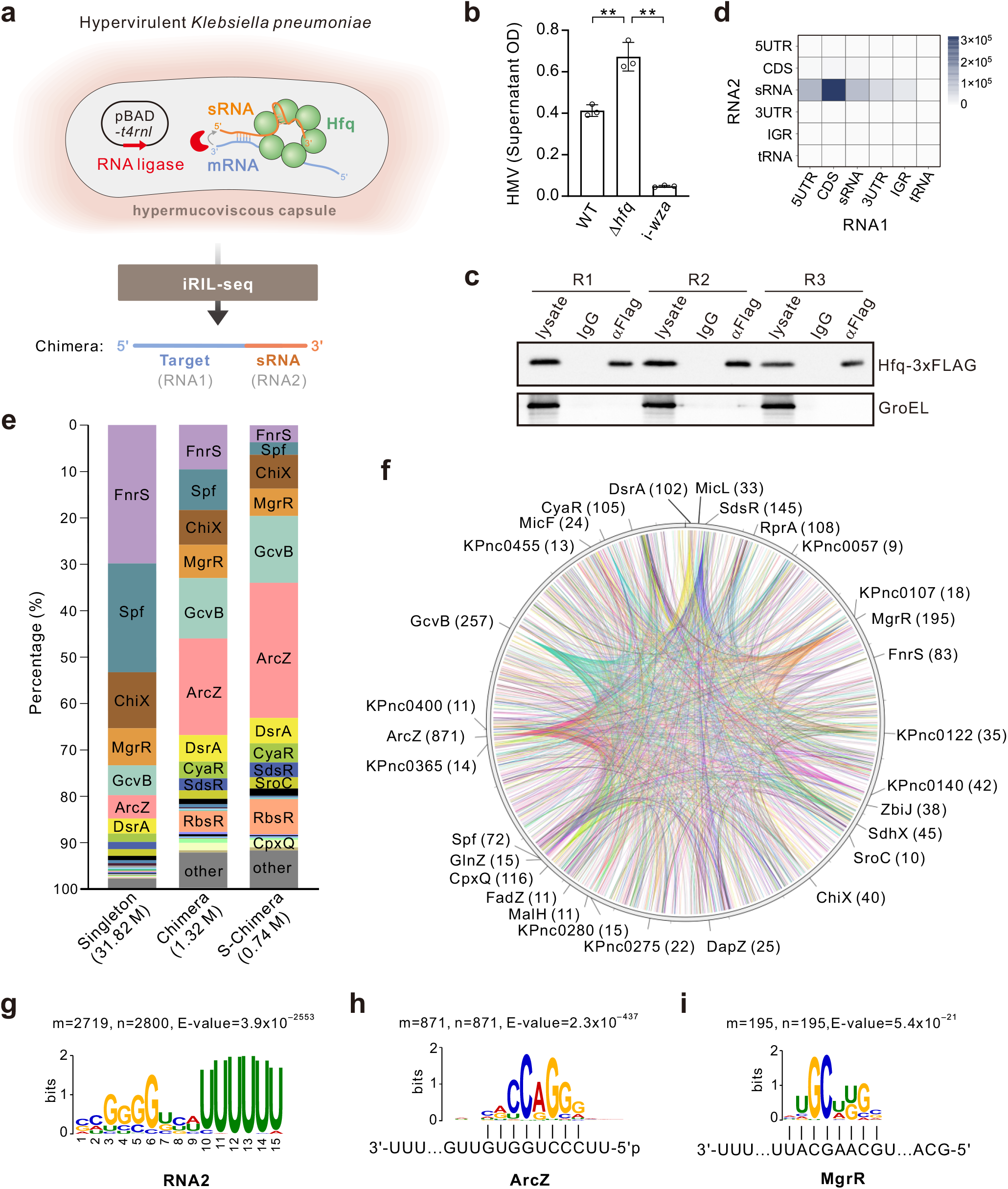
iRIL-seq analysis charts the global RNA-RNA interactome in hypervirulent *K. pneumoniae*. **(A)** Schematic of the iRIL-seq approach for *in vivo* RNA-RNA interactome profiling in HvKP. Interacting RNAs bound to Hfq are ligated by T4 RNA ligase 1, whose expression was induced briefly from pBAD-*t4rnl1* (pYC582) in live *Klebsiella* cells. Hfq-bound transcripts (singleton) and ligated fragments (chimeras) are enriched by Hfq-coIP and analyzed by deep sequencing and *in silico* analysis. **(B)** Hfq regulates hypermucoviscosity (HMV) in HvKP. Overnight cultures were centrifuged at 1000 *g* for 5 min. The OD_600_ of the supernatant was determined and normalized to the OD_600_ of starting culture before centrifugation. The acapsular i-*wza* mutant served as a (non-mucoid) negative control. N=3 biological replicates, bars indicate mean ± SD. *, *p* <0.05; **, *p* <0.01; Student’s *t*-test. **(C)** Western blot confirming IP of Hfq. Strains carrying *hfq*-3xFLAG were grown to OD_600_∼3.0 in LB and then treated with 0.2% L-arabinose to induce the expression of T4 RNA ligase for 30 min. Lysates were subjected to coIP with anti-FLAG monoclonal antibody, or with mouse IgG antibody as a control. The lysates and bound fraction after coIP were examined by western blot analysis with anti-FLAG. GroEL served as as a loading control and negative control for coIP. **(D)** Number of transcript types in significant chimeras (S-chimeras) for each combination of genomic elements. RNA1 and RNA2 constitute the 5’ portion and 3’ portion of a chimera, respectively. The numbers of interactions are indicated for each category in brackets. **(E)** Relative abundance of sRNAs detected in singleton and chimeric fragments. The percentage was obtained by dividing the number of fragments involving a given sRNA by the total fragments from sRNAs. The sRNA annotations can be found in Supplementary Table S2. **(F)** Circos plot representation of the identified RNA-RNA interaction network in *K. pneumoniae*. Labeled sRNAs interact with at least ten putative targets, and their interactions are shown in color. The number of interactions for each sRNA is indicated in brackets. A complete list of significant interactions can be found in Table S4. **(G)** Sequence motifs identified in RNA2 in S-chimeras using MEME. m, number of target sequences containing the motif; n, total number of target sequences. **(H-I)** Representative motifs identified using MEME in RNA1 from S-chimeras for ArcZ and MgrR. Motifs shown are complementary to the cognate sRNAs. m, the number of target sequences containing the motif; n, the total number of target sequences. More motifs can be found in Figure S2.

Here, we systematically profiled the RNA-RNA interactome of HvKP using an *in vivo* RNA proximity-ligation approach (iRIL-seq, **Figure 1A**, Liu et al., 2023) to decipher the RNA-mediated regulatory network. We discovered a plethora of Hfq-associated sRNA regulators of HMV and their interacting target mRNAs, establishing an intracellular RNA-RNA interactome map of HvKP and providing molecular insights into the role of post-transcriptional control in HMV. Our study revealed that the conserved enterobacterial sRNA ArcZ and its key targets are crucial regulators of HMV and *Klebsiella* pathogenesis in mouse models. ArcZ is transcriptionally activated by the master regulator of catabolite repression CRP, establishing a new link between central metabolism and HMV in *K. pneumoniae*.

## RESULTS

### Hfq and Hfq-associated sRNAs regulate hypermucoviscosity in K. pneumoniae

Mucoid HvKP resists sedimentation by low-speed centrifugation (1000 g), and thus the OD_600_ of the supernatant from 1 OD of bacteria has been established as a quantitative readout for HMV (Dorman et al., 2018; Walker et al., 2019). Using this method, we first examined whether the major RNA chaperone Hfq, and potentially sRNAs, might regulate HMV (**Figure 1B**). Indeed, we observed a significant increase in supernatant OD values for a Δ*hfq* mutant, compared to the parental strain ATCC 43816 (KPPR1), a well-known model HvKP strain used as wild-type throughout this study. As a negative control, an i-*wza* acapsular mutant, which carries a loss-of-function insertion in the capsule biosynthesis gene *wza* (Dorman et al., 2018), was not mucoid and was pelleted completely by low-speed centrifugation. This result suggested that Hfq and Hfq-associated sRNAs may suppress HMV at the post-transcriptional level in hypervirulent *K. pneumoniae*.

### iRIL-seq profiling of the RNA-RNA interactome in K. pneumoniae

To map the Hfq-mediated RNA regulatory network in HvKP, we harnessed the iRIL-seq approach (intracellular RIL-seq) recently established in our lab (Liu et al., 2023), which enables rapid profiling of the RNA-RNA interactomes in live bacteria. Transformation of the pBAD-*t4rnl1* plasmid (pYC582) into *Klebsiella* allowed us to express T4 RNA ligase 1 for proximity-ligation of RNAs *in vivo*, followed by co-immunoprecipitation (coIP) to enrich the Hfq-bound RNA transcripts (singleton reads) and ligated RNA fragments (chimeras: sRNA-mRNA, sRNA-sRNA, etc) (**Figure 1A**). The in vivo ligation followed by Hfq-FLAG-coIP (Hfq-coIP) was carried out in triplicate in cells at early stationary phase of growth in LB (OD_600_ of 3.0), with a monoclonal anti-FLAG antibody for IP and mouse IgG as a negative control (**Figure 1C**). The resulting RNA samples were purified and subjected to Illumina paired-end sequencing (**Table S1**).

Bioinformatic analysis of the sequencing reads confirmed a strong enrichment (>10-fold) of Hfq-associated sRNAs (singleton) in the Hfq-coIP samples over the IgG controls (**Figure S1**). 83 Hfq-associated sRNAs were detected in *K. pneumoniae*, including 43 sRNAs conserved in the Enterobacteriaceae family (including *E. coli* and *Salmonella enterica*), as well as 40 novel sRNA candidates in *K. pneumoniae* (**Table S2**). Hfq also directly bound to a large number of mRNAs. ∼650 mRNAs (singleton) were enriched in our dataset > 3-fold (**Table S3**), indicating a large Hfq regulon exists in HvKP.

Ligation-derived chimeric fragments containing Hfq-associated sRNAs were also enriched by Hfq-coIP, occupying ∼50% of all chimeric reads (**Figure S1**). We selected the significant chimeras (S-chimeras), defined as those with at least 10 reads and *p_adj_*<0.05 using the Fisher’s exact test as previously reported (Liu et al., 2023; Melamed et al., 2016), for subsequent analyses. The vast majority of S-chimeras (>90%) contained an sRNA at the 3’ half of chimera (RNA2), whereas the 5’ half (RNA1) comprised mostly mRNAs and sRNA fragments, which we consider binding partners of the sRNA (**Figure 1D**). Under the condition analyzed here, eight sRNAs represented 90% of all singleton reads, whereas the ArcZ sRNA was highly enriched and ranked as the most abundant sRNA in S-chimeras (**Figure 1E**).

Our analysis of the S-chimeras identified a total of ∼2800 RNA-RNA interactions involving 76 sRNAs and 1264 target mRNAs, illustrating a large and complex RNA-RNA interaction network in HvKP (**Figure 1F**). To better understand these interactions, we systematically analyzed the S-chimeras for sequence motifs. As expected, this analysis revealed a uridine stretch in the RNA2 portion at 3’ ends, consistent with rho-independent terminators found in most sRNAs (**Figure 1G**). The RNA1 portion likely contains target mRNAs that basepair with the sRNA portion in RNA2 (Liu et al., 2023). Indeed, our analyses revealed many sequence motifs in RNA1 that show complementarity to the cognate sRNA in RNA2 (**Figure 1 H-I**). In total, we identified significant sequence motifs for 15 sRNA in S-chimeras (**Figure S2**), including several conserved enterobacterial primary (FnrS, MgrR, CyaR) and processed sRNAs (ArcZ, CpxQ, RprA), as well as a new *Klebsiella* sRNA candidate (KPnc0140). Complementary sequence motifs were discovered in over 90% of mRNA genes in RNA1 with the MAST p-value <0.001, bolstering the high reliability of target identification using iRIL-seq analysis *in vivo*.

Members of the *Klebsiella* genus have relatively large genomes (>5 Mb) with a core genome similar to that of closely-related *E. coli* and *Salmonella*. iRIL-seq analysis in HvKP recapitulated many conserved RNA-RNA interactions previously reported in *E. coli* and *Salmonella* (**Table S4**). For example, among 871 interactions involving ArcZ, we captured its orthologous target mRNAs including *tpx*, *sdaC*, and a sponge sRNA CyaR (Iosub et al., 2020; Kim and Lee, 2020; Papenfort et al., 2009). These sRNA-sRNA sponge interactions can titrate one or the other and inhibit their binding to target mRNAs (Kim and Lee, 2020). In summary, the iRIL-seq analysis has established a comprehensive map of the RNA-RNA interactome in hypervirulent *K. pneumoniae*.

### Multiple sRNAs regulate Klebsiella hypermucoviscosity

To identify potential sRNA regulators of HMV hinted at by our Δ*hfq* phenotype (**Figure 1B**), we first inspected sRNA-mediated interactions with the ∼20 genes in the capsule biosynthesis cluster (Walker et al., 2019). This revealed 21 sRNA candidates that may regulate HMV via interaction with up to 14 genes in the cluster (**Figure 2A**). Outside the capsule locus, ∼100 genes contributing to the HMV phenotype have been previously identified (**Table S5**) (Dorman et al., 2018; Mike et al., 2021; Walker et al., 2020, 2019). Comparing the significant sRNA interactions against these HMV-related genes revealed another 27 sRNA candidates that may regulate HMV via interaction with 49 HMV-related genes (**Figure 2B**). By combing both lists, our analysis revealed a large RNA regulatory network of HMV which involves a panel of 31 sRNAs and 64 mRNAs, providing a valuable resource for the study of post-transcriptional regulation of HMV in *K. pneumoniae* (**Table S6**).

**Figure 2.**
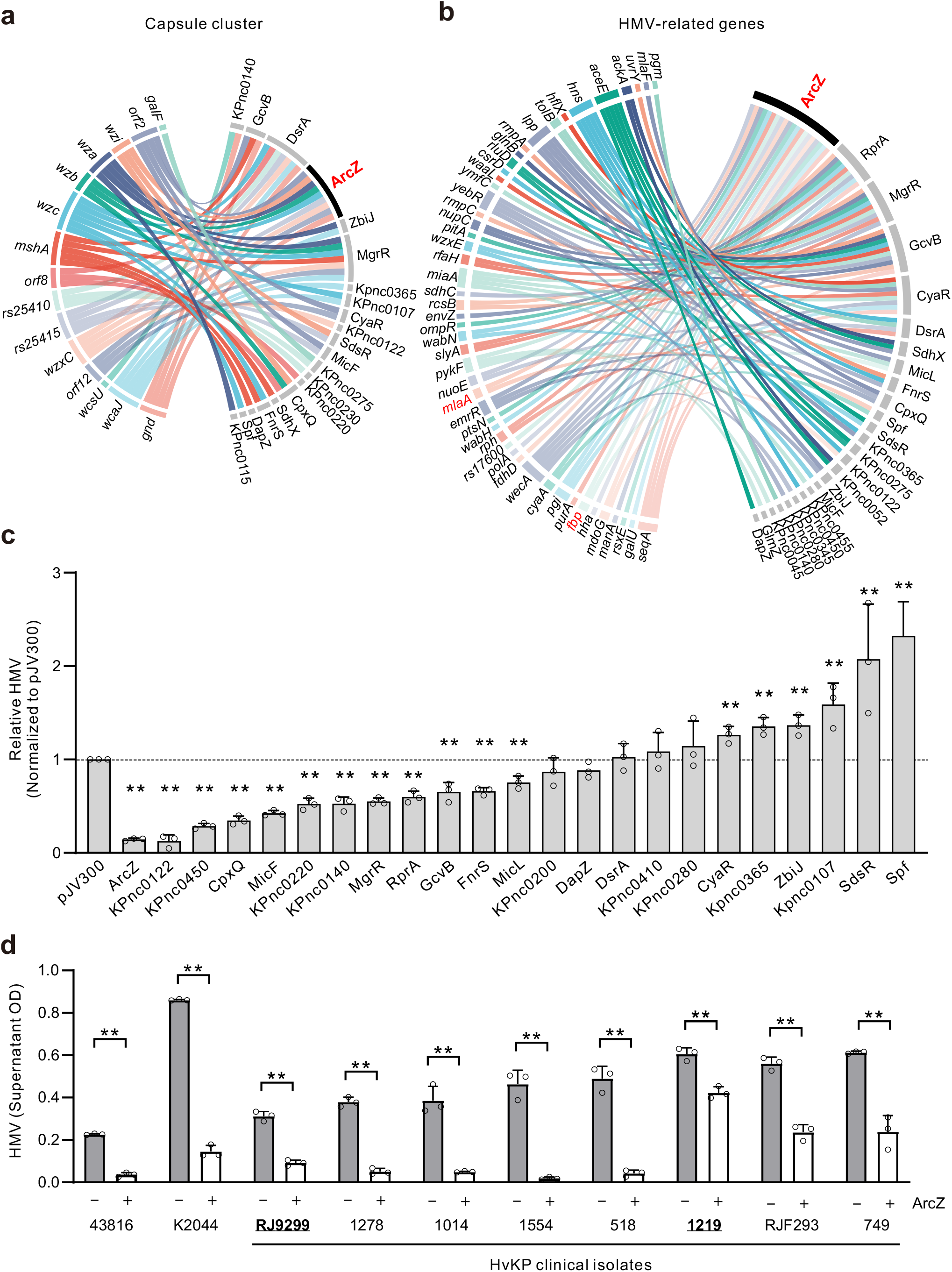
The regulatory network of hypermucoviscosity includes many sRNAs. **(A-B)** The sRNA-mRNA interaction network involving genes in the capsule biosynthesis cluster (A) and known hypermucoviscosity-regulating genes outside the capsule cluster (B). See also Figure S5 and Table S6. **(C)** The impact of sRNA overexpression on hypermucoviscosity. Strains containing sRNA expression plasmids were grown overnight in LB containing 1 mM IPTG. Cultures were centrifuged at 1,000 *g* for 5 min, and OD_600_ of supernatant was measured and normalized to the OD_600_ of the culture before centrifugation. Relative HMV values are shown relative to the basal HMV level of WT containing a pJV300 control vector. N=3 biological replicates, bars indicate mean ± SD. *, *p* <0.05; **, *p* <0.01; Student’s *t*-test. **(D)** The ArcZ sRNA reduces HMV in different model and clinical strains. Shown in bold and unlined are two clinical CR-HvKP isolates RJ9299 and 1219 (**Table S7**). N=3, bars indicate mean ± SD. *, *p* <0.05; **, *p* <0.01; Student’s *t*-test.

To experimentally validate these potential regulators of HMV, we cloned 23 sRNAs from the above list into the pZE12 vector and examined the impact of their overexpression on HMV in WT *Klebsiella*. While the pZE12 system is widely used to overexpress sRNAs in various bacteria (Corcoran et al., 2012), we found that it failed to overexpress ArcZ in *Klebsiella* (**Figure S3**). This is likely due to transcriptional repression of the P*_lacO_* promoter by ∼20 endogenous LacI-like repressors in the *Klebsiella* genome (**Figure S4**), and was overcome by a supplementation of the culture with 1 mM IPTG (**Figure S3**). Upon IPTG induction, most of the tested sRNAs showed a strong impact on HMV (**Figure 2C**), corroborating the robustness of iRIL-seq analysis (F. Liu et al., 2023). More than half (12/23) of the sRNAs tested reduced HMV vs. overexpression of a scrambled RNA in the control plasmid pJV300, whereas six sRNAs enhanced HMV to various degrees. ArcZ showed the strongest repression (∼ 6-fold) of HMV in the presence of IPTG. Interestingly, HMV was also reduced by leaky expression of ArcZ in the absence of IPTG (**Figure S3**), suggesting that HMV is highly sensitive to ArcZ levels.

Clinical *K. pneumoniae* strains are known for their extraordinary genetic diversity involving numerous sequence types and plasmid constituents (Wyres et al., 2020). To determine whether ArcZ repression of HMV is a conserved regulation or specific to our HvKP strain, we overexpressed ArcZ in a set of different HvKP clinical isolates (**Table S7**). The results in **Figure 2D** show that ArcZ induction significantly reduced HMV in all tested HvKP strains. These strains include another widely-used model strain NTUH-K2044, and a recent CR-HvKP clinical isolate (RJ9299) that is responsible for a local hospital outbreak in Shanghai, China (Zhu et al., 2022). These data strongly support that the regulation of HMV by ArcZ is a conserved mechanism among HvKP strains with diverse genetic backgrounds.

### ArcZ is a conserved repressor of hypermucoviscosity

ArcZ was initially identified in *Salmonella* in the intergenic region between *arcB* and *elbB* (a.k.a. *yhbL*), and obtained its current name due to its partial overlap with *arcB* (Papenfort et al., 2009). In *E. coli* and *Salmonella*, ArcZ is transcribed as a 129-nt long precursor sRNA (Pre-ArcZ) and further processed into the mature ArcZ sRNA (Chao et al., 2017). In *K. pneumoniae,* the *arcZ* gene is located in the same intergenic region (**Figure 3A**), and shares a conserved seed sequence with the other enterobacterial homologs (**Figure 3B**). The primary sequence of Pre-ArcZ is identical in multiple *Klebsiella* type strains. Chromosomal deletion of *arcZ* led to significantly increases in HMV (**Figure 3C**) and in the capsule amount (**Figure 3D**). These increases in Δ*arcZ* were successfully complemented by introducing a single-copy of *arcZ* gene back in the chromosome at another locus. Therefore, the endogenous ArcZ sRNA represses the capsular HMV in *Klebsiella*, and likely acts *in trans* independent of its neighboring *arcB* gene (**Figure S5**).

**Figure 3.**
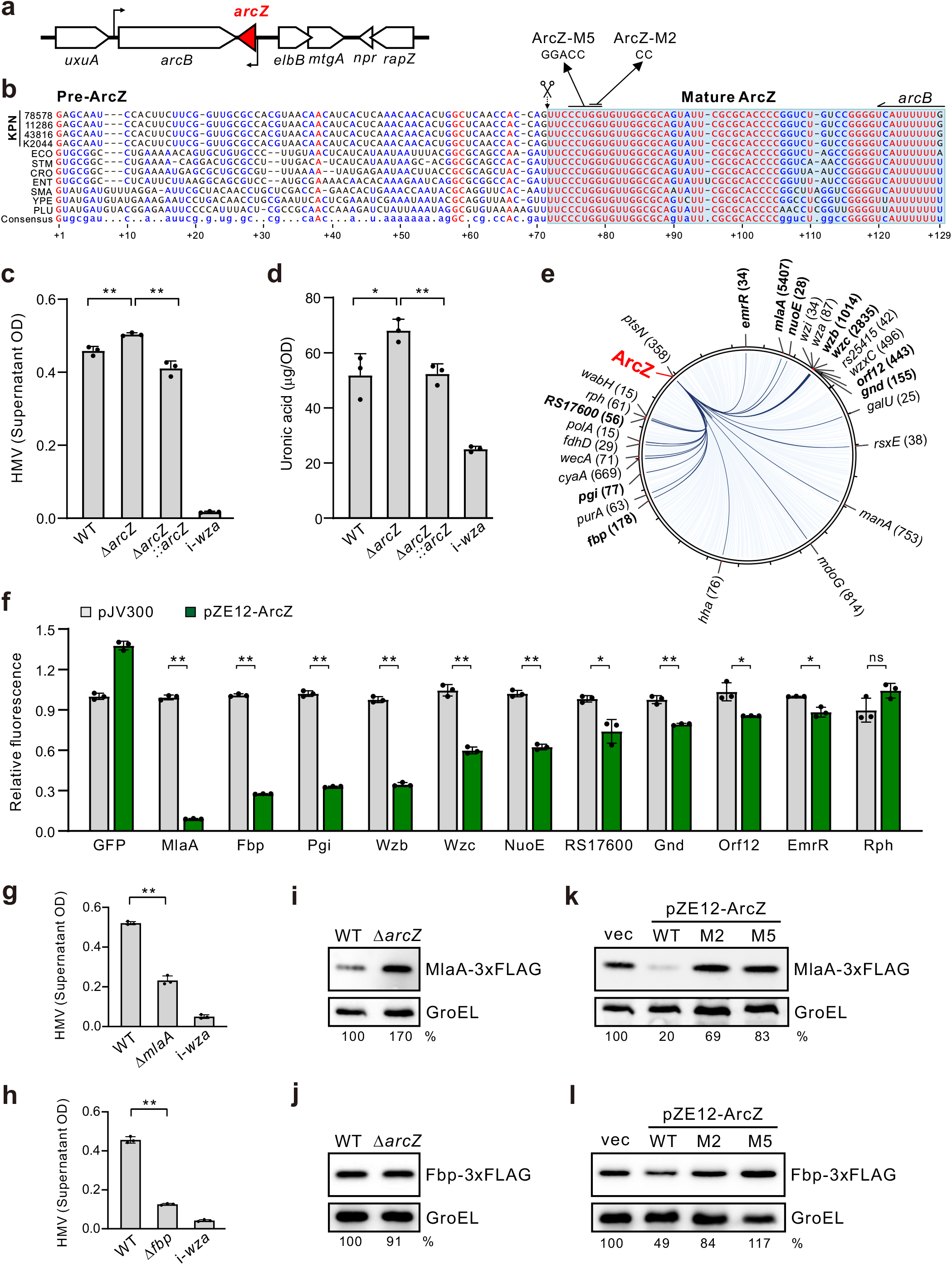
The ArcZ sRNA is a conserved regulator of multiple HMV-related genes. **(A)** Genomic context of the *arcZ* gene in *K. pneumoniae*. **(B)** Multiple sequence alignment of ArcZ sRNA. KPN, *K. pneumoniae*; ECO, *E. coli*; STM, *S.* Typhimurium; CRO, *Citrobacter rodentium*; ENT, *Enterobacter* spp.; SMA, *Serratia marcescens*; YPE, *Yersinia pestis*; PLU, *Photorhabdus luminescens*. The scissors indicate RNase E cleavage site and the start of mature ArcZ. **(C)** HMV is regulated by Δ*arcZ* and a chromosomal complementation of ArcZ. A copy of *arcZ* gene with its own promoter was inserted into *putA* locus by homologous recombination. The i-*wza* mutant serves as negative control. N=3 biological replicates, bars indicate mean ± SD. *: *p*<0.05, **: *p*<0.01; Student’s *t*-test. **(D)** ArcZ inhibits the production of capsular polysaccharides. The i-*wza* mutant serves as the acapsular control sample. N=3 biological replicates, bars indicate mean ± SD. *: *p*<0.05, **: *p*<0.01; Student’s *t*-test. **(E)** Circos plot highlighting the ArcZ interactome. Interactions involving known HMV-and capsule-regulating genes are highlighted in dark blue. The number of S-chimeras detected by iRIL-seq are indicated for each interaction in brackets. **(F)** Regulation of putative target genes by ArcZ. Fluorescence was determined in cells containing translational sfGFP reporters. The fluorescence of pJV300 (for each reporter fusion) was set to 1. N=3 biological replicates, bars indicate mean ± SD. *: *p*<0.05, **: *p*<0.01; Student’s *t*-test. **(G-H)** HMV assay for the *mlaA* (F) and *fbp* (G) mutants. The acapsular i-*wza* mutant served as non-mucoid negative control. N=3 biological replicates, bars indicate mean ± SD. *: *p*<0.05, **: *p*<0.01; Student’s *t*-test. **(I-J)** Western blot analysis of MlaA-3xFLAG (H) and Fbp-3xFLAG (I) protein levels, respectively. GroEL served as a loading control. **(K-L)** Western blot analysis of MlaA and Fbp protein levels upon ArcZ overexpression. Cultures were supplemented with 1 mM IPTG to induce ArcZ overexpression. ArcZ containing 2 nt (M2) or 5 nt (M5) mutations (panel A) were included.

According to our iRIL-seq data, ArcZ may directly interact with many mRNAs including several encoded in the capsule cluster or related to HMV (**Figure 3E**, **Figure S6**). To understand how ArcZ regulates the HMV phenotype, we selected several of these relevant genes and constructed translational GFP reporter fusions using the two-plasmid system (Corcoran et al., 2012). Of eleven different target genes, ten (91%) were significantly repressed upon ArcZ overexpression (*p*<0.05, Student’s *t*-test, **Figure 3F**). The strongest suppression was observed for *mlaA* and *fbp*, two HMV-related genes located outside of the capsule cluster. MlaA is an outer-membrane lipoprotein that functions in the retrograde phospholipid trafficking pathway (Abellón-Ruiz et al., 2017; Guest et al., 2023). The *fbp* gene encodes fructose-1,6-bisphosphatase of the gluconeogenesis pathway (Kelley-Loughnane et al., 2002). Despite their involvement in different pathways, deletion of *mlaA* and *fbp* caused a significant reduction in HMV (**Figure 3G, H**), mirroring the effect of ArcZ overexpression in *Klebsiella* (**Figure 2 C-D**).

To examine the regulation of MlaA and Fbp proteins in *Klebsiella*, we inserted chromosomal 3xFLAG tags at their respective C-termini and analyzed their levels by Western blotting. A clear upregulation (∼1.7-fold) was observed for the MlaA-3xFLAG protein when *arcZ* was deleted (**Figure 3I**), with little change for Fbp-3xFLAG (**Figure 3J**). Overexpression of ArcZ strongly reduced both levels of MlaA and Fbp proteins (**Figure 3 K-L**). This inhibition was abolished by introducing 2-or 5-nt mutations into the conserved seed sequence of ArcZ (M2 and M5, **Figure 3B**), suggesting a direct regulation involving the seed region of ArcZ.

### ArcZ represses *mlaA* and *fbp* by direct basepairing interactions

ArcZ regulates the expression of *mlaA* and *fbp* likely via base-pairing with the mRNAs. Analysis of the iRIL-seq data revealed a large number of ArcZ chimeras mapped to the 5’UTRs of both *mlaA* and *fbp* mRNAs (**Figure 4 A-B**), indicating that ArcZ basepairs at their 5’UTRs. Indeed, basepairing interactions was predicted, with both IntaRNA and RNAhybrid tools, between the conserved seed of ArcZ and the translation initiation region of both mRNAs (**Figure 4 C-D**). Using translational GFP reporters, we confirmed regulation of both targets at the post-transcriptional level. GFP fluorescence analysis showed that the target regulation requires the conserved seed region of ArcZ (**Figure 4 E-F**), consistent with Western blot results (**Figure S8**). Finally, introducing compensatory basepairing mutations in the target mRNAs (**Figure 4 C-D**), restored regulation by the ArcZ-M2 sRNA (**Figure 4 E-F**). These results show that ArcZ inhibits the translation of *mlaA* and *fbp* via direct RNA-RNA basepairing interactions.

**Figure 4.**
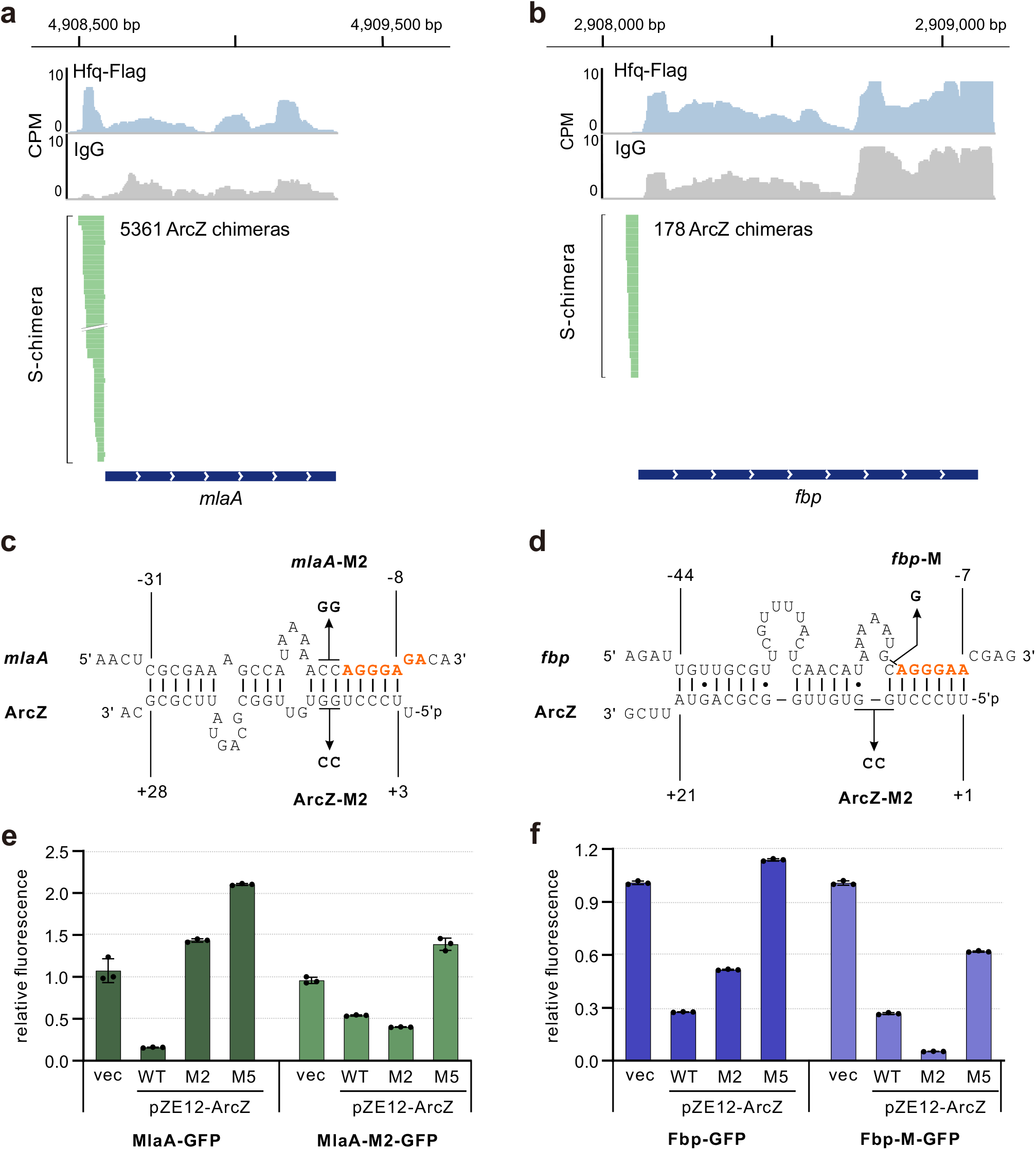
ArcZ represses *mlaA* and *fbp* via direct basepairing. **(A-B)** Browser images showing the stacked chimeric ArcZ reads mapped in the *mlaA* (A) or *fbp* (B) loci. The y-axis shows the normalized reads (CPM: counts per million); S-chimera track: Fragments of *mlaA* or *fbp* mRNAs detected in ArcZ S-chimeras in Hfq-Flag libraries. **(C-D)** Predicted base pairing for ArcZ with *mlaA* (C) and *fbp* (D). Shine-Dalgarno sequences are shown in red. Numbers above indicate the nucleotide position relative to the start codon of the respective mRNA and numbers below indicate the nucleotide position in ArcZ. The introduced mutations are indicated. **(E-F)** Compensatory basepair exchange analysis to confirm direct interactions with *mlaA* (E) and *fbp* (F). Fluorescence levels were analyzed for strains expressing MlaA-GFP and Fbp-GFP reporters.

### ArcZ and MlaA are crucial for Klebsiella virulence

Given that HMV is a major virulence factor for *Klebsiella*, we next analyzed the role of ArcZ in virulence regulation using a murine infection model (Palacios et al., 2018). Compared to the WT strain, mice infected with the Δ*arcZ* mutant showed a trend of higher CFUs recovered from lung and spleen 48 h post-infection (**Figure S9**), without reaching statistical significance. Using a more sensitive competition assay with WT and Δ*arcZ* mixed together, we observed that Δ*arcZ* showed a significant competitive advantage over WT in mice (**Figure 5A**), supporting that ArcZ is a repressor of *Klebsiella* virulence. In line with this observation, ArcZ expressed from its own promoter on multicopy plasmid (pZE12-*P_own_*-*arcZ*) led to a significant reduction of *Klebsiella* load in lungs (**Figure S9**). Constitutive overexpression of ArcZ from a stronger 16S rRNA promoter drastically reduced bacterial burden in mice by several orders of magnitude (**Figure 5 B-D**, **Figure S10**). By stark contrast, no effect on bacterial load was observed when the ArcZ-M5 mutant was overexpressed, supporting that ArcZ is a potent and specific inhibitor of HvKP pathogenesis.

**Figure 5.**
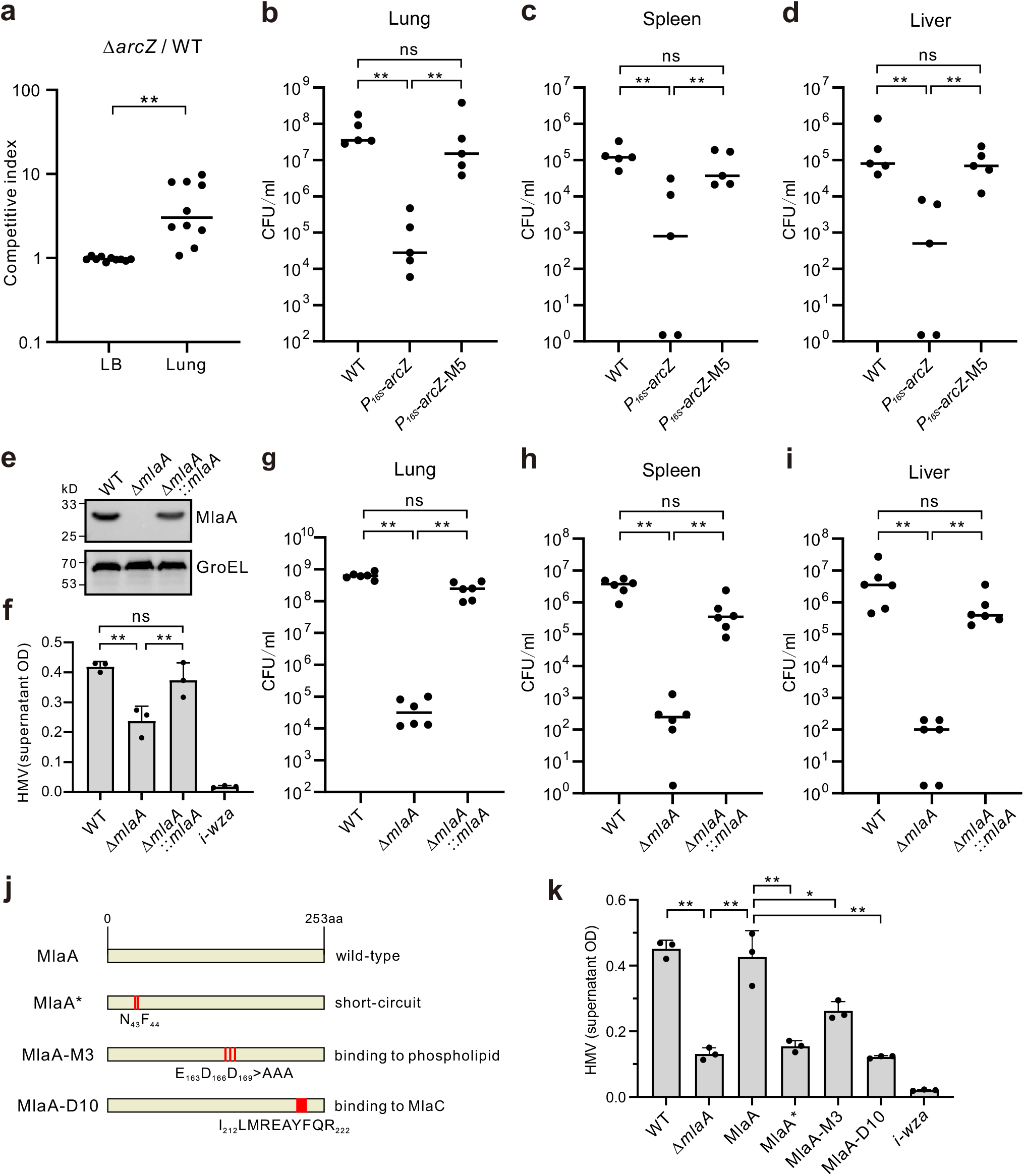
ArcZ and MlaA are required for *Klebsiella* virulence in a mouse model. (A) Competition assay using a 1:1 mixture of WT and Δ*arcZ* mutant. 6–8-week-old BALB/c mice were inoculated with 10^3^ CFU of the mixed bacteria intranasally, and scarified 48 hpi for bacterial enumeration. Growth competition in LB serves as an *in vitro* control. N=10 biological replicates, lines indicate median values. **, *p* < 0.01; Mann-Whitney test. (B-D) ArcZ overexpression significantly reduced *Klebsiella* pathogenesis in mice. Bacterial burden in different organs such as lung (B), spleen (C) and liver(D) were enumerated 48 hpi. Samples below limit of detection (100 CFU) was drawn on X-axis. N=5 biological replicates, lines indicate median value. *, *p* < 0.05; **, *p* < 0.01; ns: not significant, Mann-Whitney test. (E) Western blot verification of MlaA expression in the chromosomal deletion and complementation strains. 3xFLAG tag was introduced into the C-terminus of MlaA and used for detection by the anti-FLAG antibody. GroEL serves as a loading control. (F) MlaA is required for HMV. The HMV level was determined with overnight cultures of the indicated strains. Chromosomal complementation was constructed by inserting a copy of *mlaA* gene with its own promoter into *putA* locus using homologous recombination. The acapsular i-*wza* mutant served as negative control. N=3 biological replicates, bars indicate mean ± SD. *, p <0.05; **, p <0.01; Student’s *t*-test. (G-I) MlaA is required for *Klebsiella* pathogenesis. Mice were infected with 10^3^ CFU of the indicated strains via intranasal instillation. Bacterial burdens in lung, spleen and liver were analyzed 48 h pi. Lines indicate the median bacterial burden in each organ. N=6 biological replicates, lines indicate median value. *, p < 0.05; **, p < 0.01; ns: not significant, Mann-Whitney test. (J) Mutations introduced in the MlaA protein variants and their associated molecular functions. (K) HMV analysis for strains with chromosomal MlaA variants. The *mlaA* gene containing desired mutations was introduced back into the chromosome of Δ*mlaA* strain. N=3 biological replicates, bars indicate mean ± SD. *, p <0.05; **, p <0.01; Student’s *t*-test.

The reduced virulence may be due to the diminished expression of ArcZ target genes. Our initial competition experiments revealed that Δ*mlaA* has reduced fitness in mice. The *mlaA* mutant was significantly outcompeted in lungs, whereas it did not show any fitness defect in LB *in vitro* (**Figure S10**). When infecting mice with either WT or Δ*mlaA* in separation, we confirmed that Δ*mlaA* is severely attenuated (**Figure 5 G-I**), generating ∼1000-fold fewer CFU recovered from lung, spleen, and liver 48 h p.i. compared to WT (*p*<0.001, Mann-Whitney test). Next, we complemented the Δ*mlaA* mutant by introducing a single copy of *mlaA* gene back to the chromosome at another locus. This complementation strain not only restored the expression of MlaA protein (**Figure 5E**), and also successfully rescued HMV and pathogenesis similar to the WT levels (**Figure 5 G-I**). Collectively, these data show that the ArcZ sRNA and its target *mlaA* are crucial regulators of HMV and *Klebsiella* virulence in mice.

MlaA, together with the MlaBCDEF proteins, constitute a phospholipid retrograde transport machinery that maintains lipid asymmetry of the outer-membrane (Abellón-Ruiz et al., 2017; Malinverni and Silhavy, 2009; Sutterlin et al., 2016). MlaA removes misplaced phospholipids in the outer leaflet of outer-membrane, and transports them back to the inner membrane via periplasmic MlaC and the MlaBDEF membrane channel. To better understand the mechanism how MlaA contributes to HMV, we constructed a series of MlaA variants based on the recently solved Cryo-EM structures (Abellón-Ruiz et al., 2017; Tang et al., 2021; Yeow et al., 2023), generating a number of single-copy chromosomal *mlaA* mutant strains (**Figure 5J**). Subsequent analysis revealed that the HMV regulation by MlaA is dependent on several active sites, including residues E163/D166/D169 that are required for phospholipid binding (Abellón-Ruiz et al., 2017), and residues I212-R222 required for interaction with MlaC (Yeow et al., 2023). These data indicate that disruption of phospholipid retrograde transportation is crucial for HMV, rather than some unknown activity of the MlaA protein in *Klebsiella*. The MlaA* mutant, which induces a ‘short-circuit’ of phospholipid flux in the outer-membrane (Abellón-Ruiz et al., 2017; Sutterlin et al., 2016), also led to reduced HMV levels, indicating the homeostasis of phospholipid is crucial for HMV. Altogether, our genetic characterizations suggest a molecular model in which MlaA removes free phospholipids in the outer leaflet of outer membrane, provides sufficient space to permit the retention of capsular polysaccharide chains to the cell surface, thus promoting the HMV phenotype (**Figure 6E**).

**Figure 6.**
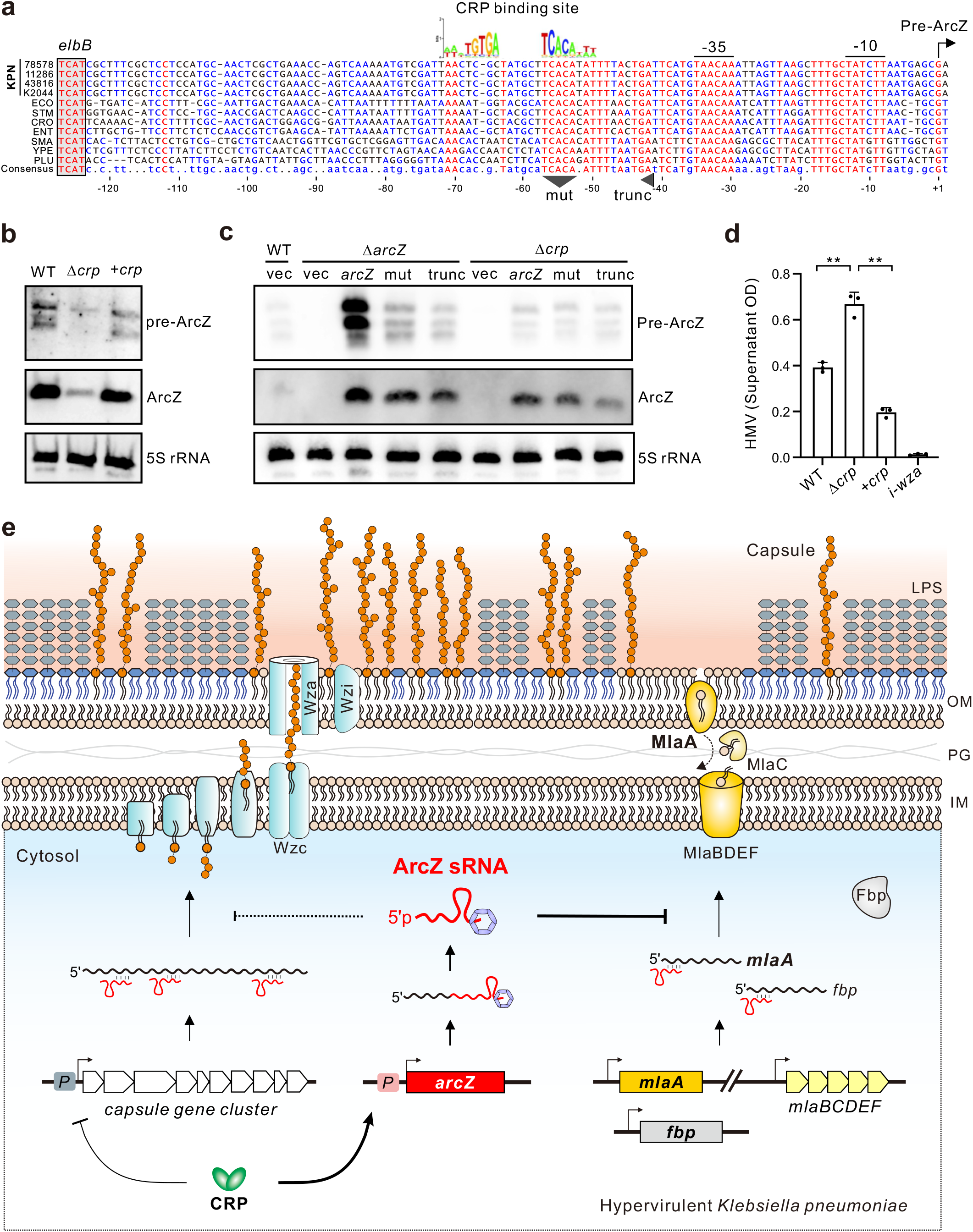
CRP activates the transcription of ArcZ in *K. pneumoniae*. **(A)** Multiple sequence alignment of the ArcZ promoter region. The putative CRP-binding site is labeled. **(B)** Expression of ArcZ requires the transcriptional regulator CRP. Northern blot was performed using 10 mg total RNA extracted from the indicated strains. Pre-ArcZ was detected using probe YCO-1838, the mature ArcZ was detected using probe YCO-1745. 5S rRNA served as a loading control. **(C)** Northern blot analysis to identify critical region in *arcZ* promoter. ArcZ expression from pZE12-based plasmids carrying different *arcZ* genomic regions. Vec, empty vector control (pJV300); *arcZ*, plasmid carrying a *arcZ* gene with native promoter; Mut: mutation of the conserved TCACA sequence. Trunc: truncation of the ArcZ promoter from -41 bp upwards without disturbing the *arcZ* gene. 5S rRNA was probed as a loading control. **(D)** CRP negatively regulates HMV. Overnight cultures of the indicated strains were centrifuged at 1000 *g* for 5 min. N=3 biological replicates, bars indicate mean ± SD. *, *p* < 0.05; **, *p* < 0.01; Student’s *t*-test. **(E)** Regulatory circuit and model for ArcZ-mediated HMV regulation in *K. pneumoniae*. Mucoid capsular polysaccharides are synthesized and translocated onto outer-membrane (OM) dependent on protein products translated from the capsule gene cluster. The transcription of capsule genes is repressed by CRP via direct binding to the promoters; whereas their translation may be repressed by ArcZ at the RNA level. CRP also activates the transcription of ArcZ precursor, which is further processed into a shorter mature ArcZ sRNA. Via direct RNA base-pairing interactions, ArcZ binds to the HMV-related mRNAs such as *mlaA* and *fbp* and inhibit their translation initiation. Reduced MlaA synthesis disrupts OM lipid asymmetry and homeostasis of phospholipid retrograde transport from outer leaflet, thereby interfering with the retention of capsular polysaccharides on the OM and proper level of HMV as well as virulence in mice. OM: outer-membrane. IM: inner-membrane. PG: peptidoglycan. LPS: Lipopolysaccharides.

### Transcriptional activation of ArcZ by CRP

To better understand the regulatory circuit of ArcZ, we sought to investigate how the sRNA is activated in *K. pneumoniae*. Previous studies in *E. coli* and *Salmonella* have shown that ArcZ is repressed by the ArcAB two-component system under low-oxygen conditions (Mandin and Gottesman, 2010; Papenfort et al., 2009), but it is unknow which upstream signals activate ArcZ expression in *K. pneumoniae*. Interestingly, multiple sequence alignment of the ArcZ promoter region from *K. pneumoniae* and other enterobacteria revealed a conserved binding site potentially recognized by CRP, the master regulator of catabolite repression (**Figure 6A**). In line with this observation, induction of catabolite repression by glucose suppressed ArcZ expression levels, suggesting that CRP activates ArcZ transcription (**Figure S11**). We next constructed a Δ*crp* mutant and observed a strong reduction in ArcZ levels. This reduction in Δ*crp* was fully restored to WT levels by the *crp* gene provided *in trans* (**Figure 6B**). Furthermore, the *arcZ* gene with its own promoter (pZE12-*P_own_*-*arcZ*) drove higher levels of Pre-ArcZ only in strains with *crp* but not in the Δ*crp* mutant (**Figure 6C**). Truncating the potential CRP-binding region from -41 bp upstream (“trunc, Figure 6A; without disturbing the -35 and -10 boxes), or mutating the conserved TCACA motif, both reduced the levels of ArcZ (**Figure 6C**). Although a molecular interaction remains to be proven, these results strongly argue that CRP directly activates the ArcZ promoter. CRP itself acts to repress HMV in *K. pneumoniae*. Deletion of *crp* significantly enhanced the mucoid phenotype, which was fully complemented by the *crp* gene *in trans* (**Figure 6D**). Therefore, we have identified a novel regulatory circuit of HMV, in which the master regulator of catabolite repression CRP activates the ArcZ sRNA, both acting to suppress HMV in *K. pneumoniae* at the transcriptional and post-transcription level, respectively (**Figure S12**, **Figure 6E**).

## DISCUSSION

Post-transcriptional control at the RNA level is crucial for bacterial pathogens to coordinate virulence and fitness under changing conditions during host infection. In this study, we generated a comprehensive RNA-RNA interactome map in the hypervirulent pathotype of *K. pneumoniae*, revealing the post-transcriptional regulatory network of virulence control in a superbug of global concern. We discovered that HMV in HvKP is regulated by the major RNA chaperone Hfq and a large number of Hfq-associated sRNAs. Our study further demonstrated the molecular mechanism how ArcZ, as an Hfq-dependent sRNA, regulates HMV and virulence via direct sRNA-mRNA interactions, adding support to the evolving theory that HMV is critically regulated in *K. pneumoniae* during the process of host infection (Chu et al., 2023; Ernst et al., 2020; Khadka et al., 2023). Our successful identification of key HMV regulators and regulatory circuit in this study may illustrate a new paradigm of using RNA-RNA interactome to guide functional RNA screens with mechanistic precision to understand important traits in different organisms.

Our systems-wide profiling of the RNA interactome in *K. pneumoniae* was achieved by using the iRIL-seq approach, which was recently developed and established in *Salmonella* in our lab (Liu et al, 2023). Taking advantage of expressing T4 RNA ligase *in vivo* (Han et al., 2016) and Hfq-coIP (Chao et al., 2012), iRIL-seq enables proximity-ligation between interacting RNAs in live bacteria and enriches the ligated RNA fragments bound to Hfq. Our *in vivo* approach is highly streamlined compared to crosslinking-based approaches RIL-seq and CLASH (Iosub et al., 2020; Melamed et al., 2018), eliminating the laborious *in vitro* ligation and digestion steps. iRIL-seq has been proven to effectively capture sRNA-target interactions at the genome-wide scale in *Salmonella* with high accuracy (Liu et al, 2023). Here the RNA-interactome data obtained in *K. pneumoniae* further support the utility of iRIL-seq analysis. Consensus sequence motifs complementary to sRNAs were identified in over 90% of target mRNA genes. Given its simple setup and streamlined workflow, our HvKP dataset further showcase iRIL-seq as a powerful generic approach that can be easily applied to many different organisms to chart functional RNA regulatory networks.

Our study has discovered a previously unknown large RNA regulatory network involving many HMV-related genes in the HvKP RNA interactome, while numerous known sRNA-mRNA and sRNA-sRNA interactions from *E. coli* and *Salmonella* were also recapitulated. The sRNA regulators of HMV include many core-genome encoded sRNAs conserved in enterobacteria, and also new sRNA candidates in *Klebsiella* (**Figure 2C**, **Table S2**). Several of these sRNAs have been previously studied in model organisms. CpxQ and MicF function in the envelope stress pathway to target membrane proteins (Chao and Vogel, 2016; Corcoran et al., 2012); MgrR in the PhoPQ pathway regulates LPS modification and magnesium transport (Moon and Gottesman, 2009; Yin et al., 2019); and RprA is an activator of RpoS involved in general stress responses (Sedlyarova et al., 2016). These sRNAs with diverse functions all strongly inhibit HMV in *K. pneumoniae*, suggesting that sRNAs from several distinct pathways might converge to modulate *Klebsiella* virulence during infection. *Klebsiella* sRNA candidates such as KPnc0122, KPnc0450 and KPnc0107 also repressed HMV, providing opportunities for future molecular and functional characterizations.

Our study discovered that the enterobacterial core sRNA ArcZ is integrated into a new regulatory circuit and is endowed with a new function in *K. pneumoniae* (**Figure 6E**). In *E. coli* and *Salmonella,* the promoter of ArcZ has been shown to be repressed by the ArcAB two-component system (Mandin and Gottesman, 2010; Papenfort et al., 2009). Our results suggest that ArcZ, instead of being repressed by ArcAB (**Figure S8**), is activated by the catabolite repression protein CRP in *K. pneumoniae*. This indicates that ArcZ might have evolved with new regulatory functions under the control of CRP. Functional enrichment analysis for the ArcZ targets revealed several significant pathways related to carbohydrate and polysaccharide metabolism (**Figure S6A**), consistent with the observed repressive roles on HMV (**Figure 2 C-D**) and capsule production (**Figure 3D**, **S3C**). Total RNA-seq analysis also revealed significant down-regulation of capsule genes upon ArcZ overexpression (**Figure S7**), although a direct regulation remains to be established. Our results suggest that even conserved sRNAs may have evolved distinct functions in different organisms. For example, a recent study showed that the homologs of ArcZ in *Photorhabdus*/*Xenorhabdus* regulate metabolite production and symbiosis with nematodes (Neubacher et al., 2020).

CRP controls one of the largest regulons to modulate bacterial carbon metabolism and energy expenditure. Consistent with previous reports (Lin et al., 2013, 2018), our study suggests that CRP is a negative regulator of HMV. CRP was shown to repress the transcription of capsule gene clusters via direct binding to the promoters. In the meantime, CRP activates the ArcZ sRNA, which in turn binds and represses many other HMV-related mRNAs. ArcZ may function as an extension of CRP to repress more HMV genes at the post-transcriptional level (**Figure 6E**). We note that the CRP pathway may be more complicated than this simple model. Besides ArcZ, CRP regulates two other sRNAs, Spf and CyaR, according to studies in *E. coli* and *Salmonella* (Beisel and Storz, 2011; Papenfort et al., 2008). Both sRNAs positively regulated HMV, in contrast to the repressive roles of ArcZ and Hfq. Thus, sRNAs within the CRP pathway both negatively and positively regulate HMV, indicating the existence of complicated regulatory loops fine-tuning HMV at the post-transcriptional level in *K. pneumoniae*.

We show for the first time that the phospholipid trafficking pathway is crucial for *Klebsiella* virulence and that MlaA is an essential virulence factor in HvKP. As a key target of ArcZ, *mlaA* encodes the crucial outer membrane component of the Mla phospholipid retrograde transportation system that maintains lipid asymmetry and membrane integrity (Abellón-Ruiz et al., 2017; Guest et al., 2023; Malinverni and Silhavy, 2009). Deletion of *mlaA*, without showing any growth or fitness defect in broth, resulted in a strong reduction of HMV and a severe attenuation of virulence in the murine model of infection. Transposon disruption of *mlaBCDEF* genes also led to decreased HMV (Dorman et al., 2018), supporting the involvement of the entire Mla system and its molecular function. Our further mutational analysis of MlaA revealed that phospholipid transport activity is crucial for capsule retention and HMV, providing a new mechanism underlining the HMV phenotype. Because the Δ*mlaA* mutant still retains a low level of HMV higher than the acapsular control, its strong attenuation in virulence makes Δ*mlaA* an attractive candidate to develop live attenuated vaccine. MlaA might also represent an interesting drug target located on the bacterial surface for the future development of therapeutics against HvKP. On the same note, ArcZ may also be exploited as a potential anti-infection drug in the form of antisense RNA therapeutics (Popella et al., 2021; Vogel, 2020). Our data demonstrated that ArcZ is effective to reduce HMV in a variety of clinical HvKP strains including CR-HvKP isolates, which urgently demand new treatment options in clinics.

## MATERIAL AND METHODS

### Bacterial strains and culture conditions

Hypervirulent *Klebsiella pneumoniae* ATCC 43816 is used as wild-type. Mutant strains generated in this study are described in **Supplementary Table S8**. Unless otherwise stated, bacteria were grown in LB medium (10 g/L tryptone, 5 g/L yeast extract, and 5 g/L NaCl) with 220 rpm aeration or on LB agar at 37°C. Where appropriate, antibiotics were used at the following concentrations: 20 µg/ml chloramphenicol, 50 µg/ml kanamycin, 50 µg/ml spectinomycin, 100 µg/ml hygromycin B, and 50 µg/ml apramycin sulfate. For induction of sRNA expression from pZE12 plasmids, 1 mM IPTG was used.

Deletion strains and chromosomally 3xFLAG epitope tag strains were constructed using the λ Red system (Huang et al., 2014; Uzzau et al., 2001). To construct the Δ*arcZ* mutant, wild-type *Klebsiella* cells carrying the temperature-sensitive pACBSR-Hyg helper plasmid expressing λ Red recombinase were transformed with 1 µg dsDNA fragments, which were amplified using YCO-1438/-YCO-1439 and pFLP-hyg as template. Desired recombinants were confirmed by resistance to apramycin, and by colony PCR with YCO-1038/-YCO-1039 and Sanger sequencing. To introduce 3xFLAG epitope tags, DNA fragments were amplified by PCR with primer pairs YCO-1447/1448 and pSUB11 as template, and electroporated into competent cells expressing λ Red recombinase from pACBSR-Hyg. Correct transformants were selected on kanamycin plates and confirmed by colony PCR.

Chromosomally complemented strains were constructed using the λ Red system and from pACBSR-Hyg (Huang et al., 2014; Uzzau et al., 2001). DNA fragments were amplified from pXG10 vectors containing resistance maker and *P_own_-arcZ* or *P_own_-mlaA* fragments by primer pairs YCO-2321/2322, and electroporated into mutant strains lacking either *arcZ* or *mlaA*. Point mutations in MlaA were first introduced into the pXG10-*P_own_-mlaA* vector by site-directed mutagenesis using primers described in Supplementary Table S8. Mutated *mlaA* fragments were amplified using oligos YCO-2321/2322, and electroporated into Δ*mlaA* competent cells expressing λ Red recombinase. The DNA fragments were inserted into the *putA* locus on chromosome by homologous recombination as previously described (Westermann et al., 2016). Correct transformants were selected on chloramphenicol plates, and confirmed by colony PCR and Sanger sequencing.

### Clinical strain isolation and characterization

*K. pneumoniae* clinical isolates were collected from Peking University Third Hospital **(Supplementary Table S7)**. Clinical characteristics were obtained from electronic medical records. Strains were initially identified by MALDI-TOF mass spectrometry and then confirmed via the Vitek 2 system. String-test positive *Klebsiella* isolates were further characterized by whole-genome sequencing (WGS) and analyses using the Kleborate software (Lam et al., 2021). Antibiotic sensitivity tests were performed on the six selected isolates by the Vitek 2 or disk diffusion test, according to the 2022 Clinical and Laboratory Standards Institute (CLSI) guidelines. The antimicrobials tested were cefepime (FEP), ceftazidime (CAZ), aztreonam (ATM), imipenem (IPM), meropenem (MEM), piperacillin/tazobactam (TZP), cefperazone-sulbactam (CSL), amikacin (AMK), tobramycin (TOB), colistin (COL), levofoxacin (LVX), ciprofoxacin (CIP), minocycline (MNO), tigecycline (TGC), polymyxin (POL), cefderocol (FDC), ceftazidime/avibactam (CZA) and trimethoprim/sulfamethoxazole (SXT).

### Plasmid construction

The oligonucleotides and plasmids used in this study are listed in **Supplementary Tables S9** and **S10**, respectively. To express a plasmid-borne *t4rnl1* gene from the tightly controlled, L-arabinose-inducible pBAD promoter, pBAD-Hyg was amplified by PCR with oligonucleotides YCO-0453/-0454, and the *t4rnl1* gene was amplified from pKH13-*t4rnl1* using oligonucleotides YCO-0905/-0906. Vector and insert were ligated following restriction digestion with *Xba*I. Overexpression plasmids for sRNAs were constructed according to (Corcoran et al., 2012) using pZE12-luc as the scaffold plasmid. Here, an *Xba*I-digested PCR product obtained by amplification of pZE12-luc with primers YCO-0557/-0558 was ligated to sRNAs amplified from genomic DNA, and similarly digested with *Xba*I. Construction of GFP translational fusion plasmids were carried out principally as described in (Corcoran et al., 2012), using the pXG10-SF or pXG30-SF. The pXG10-SF vector was used for monocistronic genes, the pXG30 vector for operons. Briefly, the pXG10-SF backbone was amplified with YCO-1437/-1398, and pXG30-SF backbone was amplified with YCO-1806/-1398 or YCO-1940/-1398. The 5’ regions of target genes, including regions captured in the chimeric fragments, were PCR amplified, digested with *Nsi*I/*Pst*I and *Nhe*I and cloned into pXG10-SF or pXG30-SF backbones digested with the same restriction enzymes. All the plasmids and insertions were verified by colony PCR and Sanger sequencing.

### HMV quantification

*K. pneumoniae* strains were grown overnight in 3 ml LB medium at 37°C. Next, 1 ml of culture was centrifuged for 5 min at 1,000 × *g* (room temperature), and the OD_600_ of the upper 200 μl supernatant was determined. Final readings were normalized to the OD_600_ in the input culture before centrifugation. At least three biological replicates were prepared for each strain.

### Capsule purification and quantification

*K. pneumoniae* was grown overnight in 1 mL LB medium with 220 rpm at 37 °C. Cultures were added with 100 µl the capsule extraction buffer (500 mM citric acid pH 2.0, 1% Zwittergent 3-10), incubated at 50°C for 20 min. Cells were pelleted, and the supernatant was mixed with an equal volume of isopropanol, followed by precipitation for 30 min at 4°C. The capsule was centrifuged for 10 min at maximum speed and resuspended in 150 µl PBS. Protein and membrane contaminants were removed by adding 100 µl buffer P2 (TIANGEN) and 125 µl buffer P3 (TIANGEN). The clarified supernatant was precipitated with an equal volume of isopropanol for 30 min at 4°C. The capsule was centrifuged at maximum speed for 30 min at 4°C and washed twice with 75% ethanol. The purified capsule was resuspended in nuclease-free water and confirmed on SDS-PAGE by alcian blue staining. Uronic acid assay was used to quantify the amount of capsular polysaccharides. 12.5_mM borax (Sangon) in H_2_SO_4_ was added to samples and boiled before chilling on ice. Followed by addition of 0.15% 3-hydroxydiphenol (Sigma-Aldrich), the mixture was analyzed by the absorbance at 520_nm. Uronic acid content was determined from a standard curve of glucuronic acid (Sangon) and normalized as micrograms per 1 OD of bacterial culture.

### Hfq-coIP experiments

The Hfq-coIP was performed using our previously published protocol (Chao et al., 2012). *K. pneumoniae hfq*::3xFLAG strain containing the plasmid pYC582 was grown in LB medium with 220 rpm at 37 °C. T4 RNA ligase 1 was induced for 30 min by adding 0.2% L-arabinose when the culture reached OD_600_ of 3.0. Bacterial samples (about 100 OD) were collected by centrifugation at 6,000 rpm for 20 min at 4 °C. The bacterial pellets were resuspended in 500 µl ice-cold lysis buffer (20 mM Tris pH 8.0, 150 mM KCl, 1 mM of MgCl_2_, 1 mM DTT, 0.05% Tween-20), and broke with 500 µl glass beads using the Cryolys Evolution instrument, at 7,200 rpm for 15 min at 4 °C. Lysates were collected by centrifugation at 13,000 rpm for 30 min at 4 °C, and divided into new tubes equally. Cleared lysates were respectively incubated with anti-FLAG antibody (Sigma) or IgG isolate control (Invitrogen) conjugated on the protein-G magnetic beads (Thermo-Fisher) for 1 hour at 4 °C with rotation. The beads were collected by the magnet, and washed five times with 500 µl lysis buffer and resuspended in 50 µl lysis buffer. The Hfq-bound RNA was extracted by RNA clean & concentrator columns (Sangon, Shanghai). The purified RNA was resuspended in nuclease-free water.

### RNA extraction

Total RNA was extracted using TRIzol reagent. Briefly, 4 OD of cells were collected with addition of 0.2 vol/vol of STOP solution (95% ethanol, 5% phenol), centrifuged for 5 min, 14,000 rpm at 4°C and the supernatant was discarded. For cell lysis, pellets were resuspended with 500 µl lysozyme (A610308, BBI) at 0.5 mg/ml. Cleared lysates were mixed with 1 ml TRIzol and incubated for 10 min at room temperature with shaking. Subsequently, 400 µl chloroform was added, mixed by inversion, and incubated for 5 min at room temperature to allow phase separation. After the samples were centrifuged for 15 min, 14,000 rpm at 4°C, the upper phase was transferred to a new tube and 700 µl isopropanol was added and incubated overnight at -20°C. Finally, samples were centrifuged, washed with cold 75% v/v ethanol and air-dried for 15 min. Precipitated RNA was resuspended in 30 µl nuclease-free water and stored at -80°C.

### Northern blotting

Ten micrograms of total RNA were denatured at 98°C for 2 min in RNA loading buffer (98% formamide, 1 mM EDTA, 0.1% xylene cyanol, 0.1% Bromphenol blue) and separated on 7 M urea/6% polyacrylamide gels in 1× TBE buffer. RNAs were transferred onto Hybond-N+ membranes (GE Healthcare) by electroblotting (50 V) for 1 h at 4°C and fixed to the membrane by UV crosslinking (120 mJ). Detection was performed using the Roche DIG system. Each primary probe was designed to contain an overhang sequence that is complementary the digoxigenin-labeled secondary probe YCO-0550. For instance, mature ArcZ was detected using the specific complementary probe YCO-1745, while CyaR was detected using primary probe YCO-1784. Both primary probe-hybridized sRNAs were detected with secondary probe YCO-0550. Membranes were prehybridized in DIG Easy Hyb (#11796895001, Roche) for 30 min. DNA probes were hybridized overnight at 42°C. Membranes were washed with 5× SSC/0.1% SDS, 1× SSC/0.1% SDS and 0.5× SSC/0.1% SDS buffers for 15 min each. Following one wash with maleic acid wash buffer (0.1 M maleic acid, 0.15 M NaCl, 0.3% V/V Tween-20, pH 7.5) for 15 min at 37°C and blocked with blocking solution (#11096176001, Roche) for 1 hour at 37°C, membrane was hybridized with 75 mU/ml Anti-Digoxigenin-AP (#11093274910, Roche) in blocking solution for 60 min at 37°C. Wash membrane twice (2 × 15 min) with maleic acid wash buffer and equilibrate membrane 3 min in Detection Buffer (0.1 M Tris–HCl, 0.1 M NaCl, pH 9.5). Signals were visualized by CDP-star (#11759051001, Roche) on a ChemScope imaging station and quantified using ImageJ Software.

### Western blotting

Bacterial cells were collected from cultures by centrifugation for 5 min at 12,000 rpm at 4°C, and pellets were resuspended in protein loading buffer. After heating for 5 min at 95°C, 0.05 OD_600_ equivalents of samples were separated on 12% SDS–PAGE gel and subsequently transferred to PVDF membranes (polyvinylidene fluoride, #10600023, Amersham). Membranes were blocked for 1 h with 5% (w/v) milk powder in TBS-T (Tris-buffered saline-Tween-20) and incubated overnight with primary antibody (monoclonal anti-FLAG, 1:10,000, Sigma #F1804-5MG; anti-GroEL, 1:10,000, Sigma; or anti-GFP, 1:10,000, Roche #11814460001 in 3% bovine serum albumin (BSA)/TBS-T) at 4°C. Following three washes for 10 min in 1× TBST buffer, membranes were incubated with secondary anti-mouse or anti-rabbit HRP (horseradish peroxidase)-linked antibodies (#D110087-0100 or #D110058-0100, BBI; 1:10000) diluted in 1× TBST buffer containing 1% skim milk. After another three washes for 10 min in 1× TBST buffer, chemiluminescent signals were developed using ECL Prime reagents (#C500044-0100, Sangon) visualized on a ChemScope imaging station and quantified using ImageJ software.

### iRIL-seq cDNA library construction

iRIL-seq libraries were constructed using the RNAtag-seq method as described previously (Liu et al., 2023; Melamed et al., 2018), with a few modifications. Briefly, fragmented RNA was treated with DNase I and FastAP. RNA then was ligated to the 3’ barcoded adaptor, and depleting Ribosomal RNA by Ribo-off rRNA depletion Kit (#N407-01, Vazyme). First strand cDNA was synthesized by HiScript II 1st Strand cDNA Synthesis Kit (#R211-01, Vazyme), and RNA was degraded after reverse transcription by 1 M NaOH and 0.5M acetic acid. The cDNA was ligated to a second adaptor. Every step needed to purify the sample with 2.5x Agencourt AMPure XP beads and 1.5x isopropanol. Libraries were constructed by PCR with Illumina P5 and P7 primers using Q5 High-Fidelity DNA Polymerase (#M0491L, NEB), and purified with 1.5x AMPure XP beads. The libraries were sequenced by 150 bp paired-end sequencing with an Illumina NovaSeq instrument.

### iRIL-seq data computational analysis

Data analysis was performed using the method reported by Melamed *et al*. (Melamed et al., 2018, 2016) with minor modifications. Briefly, clean reads were generated from raw sequencing reads by removing adaptor sequences and low-quality sequences using FASTXToolkit (Version 0.0.13), and reads less than 25 nt were discarded. The first 25 nucleotides of each remaining read were mapped to the genome of *K. pneumoniae* ATCC 43816 (GenBank: NZ_CP009208.1), sorting the fragments into “single” and “chimeric” fragments. Fragments that mapped within a distance of 1,000 nt were considered single, whereas fragments that mapped to two different loci were considered chimeric.

#### Finding statistically significant chimeras

To identify over-represented chimeras, we divided the genome into non-overlapping 100 nt-long windows and counted the number of sequenced fragments showing fusion between each window pair. If neighboring windows of windows appear in chimeric fragments, the size of tested windows is increased gradually in leaps of 100 nt up to 500 nt (from four windows upstream each original window to four windows downstream). We applied Fisher’s exact test to test if the chimeric fragments between two windows were statistically significantly more than expected at random based on the number of single and chimeric fragments mapped to each window. The p-values were corrected using the Bonferroni method. Chimeras with P_adj_-value < 0.05, odds ratio value > 1 and chimeric fragments > 10 were considered significant and termed S-chimeras. To detect the precise positions of ligation site for each S-chimera, chimeric fragments were then further extracted and realigned against the reference genome using BWA-MEM (parameters: -M -k10 -T10). Finally, partial alignments of segments within a single read as candidates supporting ligation point were clustering to determine the positions of ligation for each S-chimera.

#### Annotated and novel Hfq-bound sRNAs in Klebsiella

To assess whether homologs of known sRNAs are involved in Klebsiella RNA-RNA interactome, we searched by blastn the *Klebsiella* orthologs of all currently documented sRNAs in *E. coli*, *Salmonella* and orthologs of recently found sRNAs. iRIL-seq network data can be exploited to detect novel sRNAs by taking into account unique features of sRNAs in general and particular features of sRNAs included in the chimeras. Accordingly, these features include the following: (i) sRNAs have stem-loop structures followed by a poly-uracil sequence at their 3′ ends, (ii) sRNAs are enriched for Hfq, (iii) sRNAs tend to be the second RNA in the S-chimera fragment, (iv) and sRNAs are typically involved in multiple interactions with mRNA regions corresponding to CDS or 5′UTR.

#### Annotation of RNAs

The genome annotation was based on NCBI version GCF_000742755.1. The UTRs were considered to be the regions 100 nt upstream the annotated start codons and downstream of the annotated stop codon (or shorter if these regions spanned another transcript or were more likely to be a UTR of the neighboring transcript). Each region was assigned to one of the following categories: sRNA, tRNA, rRNA, hkRNA, 5’UTR, coding sequence (CDS), 3’UTR, or IGR (Intergenic region).

#### Identification of binding motifs

The target sequences of each sRNA that had at least ten targets were extracted and searched for motif using MEME (Bailey et al., 2009). To verify complementarity between the identified motifs in putative mRNA targets and the sRNA, we used MAST (Bailey et al., 2009).

#### Clustering of libraries

The reproducibility of the results was evaluated for all single and chimeric fragments. The sequenced fragments were mapped to 100 nt-long windows along the genome for each library. Each window was assigned the count of single fragments mapped to it, and then the Pearson correlation coefficient of window counts could be computed between each pair of libraries. Next, we counted the number of times each pair of windows appeared in chimeric fragments (regardless of the order of RNAs in the chimera) for each library, and calculated the Pearson correlation between libraries. Finally, we clustered the libraries by these vectors of correlation coefficients using Pheatmap.

### Total RNA-seq analysis

*K*. *pneumoniae* strains in biological triplicates were grown to OD_600_ of 3 in LB media containing IPTG (final concentration 1 mM). Cells were harvested by addition of 0.2 volumes of stop mix (95% ethanol, 5% (v/v) phenol) and snap-frozen in liquid nitrogen. RNA-seq libraries were constructed by Novogene Group, Beijing, China. The cDNA libraries were sequenced on Illumina NovaSeq X plus (150 bp paired-end). We used cutadapt 2.10 and fastx toolkit 0.0.14 to clip the adapter sequences and quality filtering. Reads unambiguously aligned to unique position in the reference genome (GenBank: NZ_CP009208.1) were preserved to calculate reads number, CPM (counts per million) and RPKM (reads per kilobase and per million) for each gene. Differential expression was tested using DESeq2, generating *p*-value and FDR based on the model of negative binomial generalized linear model.

### Fluorescence analysis

Single colonies of *Klebsiella* strains harboring superfolder GFP (sfGFP) translational fusions and sRNA expression plasmids were inoculated into LB medium and grown overnight at 37°C. The following day, 100 µl of overnight cultures were centrifuged and the pellets were resuspended in 1 ml of 1×PBS. OD_600_ and fluorescence (excitation at 476 nm and emission at 510 nm, using emission cutoff filter of 495 nm) was measured using the Agilent BioTek Synergy H1 plate reader. Fluorescence measurements were normalized to the OD_600_ of cultures. Background fluorescence from control samples not expressing fluorescent proteins was subtracted from experimental samples. The level of regulation was calculated as the ratio between the fluorescence level of a strain carrying the sRNA over-expressing plasmid and a strain carrying the control plasmid. Three biological replicates were prepared for each strain.

### Animal infections and ethics statement

All animal experiments were conducted following institutional ethics requirements under the animal facility user permit (No. IPS-26-CYJ). The infection experiments were performed according to the animal protocol (A2022053) approved by the Animal Care and Ethics Committee of Institut Pasteur of Shanghai, CAS. At least five female 6-8 weeks old Balb/C mice were used per experimental group. Mice were anaesthetized via intraperitoneal injection of tribromoethanol (0.2 ml/10 g) prior to infection. Mice were intranasally infected with approximately 10^3^ CFUs (in 25_µl PBS) per bacterial strain, or a total of 10^3^ CFUs in competition assays. Mice were weighted daily until the end of experiment. Approximately 48 h post-infection, mice were euthanized and organs including lung, liver, and spleen were processed for CFU analysis. Serial dilution of CFUs were plated on LB agar plates with appropriate antibiotics and incubated overnight at 37°C.

### Statistical analysis

All experiments were repeated at least three times. At least five mice per group were used for infection assays. Mean values and standard deviations were shown in figures. Statistical significance (*P* values) was analyzed using the GraphPad software (version 9).

## Data availability

The sequencing data have been deposited in the GEO database under accession number GSE243246. Raw data, materials and regents are available upon request.

## ACKNOWLEDGEMENTS

We are grateful to Drs. Joan Mecsas, Francesca Short, Hong-Yu Ou, Min Li, and Chao Yang for providing *Klebsiella* strains, Dr. Yunn-Hwen Gan for discussion. This study was financially supported by the National Key R&D Program of China (2022YFC2303200 and 2022YFE0111800), the Chinese Academy of Sciences (XDB0570000 and 176002GJHZ2022022MI), the Natural Science Foundation of China (32270064 and 32200031), Shanghai Municipal Science and Technology Commission (21ZR1471300 and 2019SHZDZX02), and Shanghai Super Postdoc Fellowship (R.D).

## CONFLICT OF INTEREST

We declare no conflict of interest.

## AUTHOR CONTRIBUTION

YC designed the study; KW and XL performed iRIL-seq analysis; KW characterized ArcZ and mechanisms of target regulation; XL performed capsule quantifications, characterized regulation by CRP and molecular function of MlaA; YL analyzed clinical isolates and performed mice infection experiments with help from RD and HJ; JZ, NS contributed materials and regents; KW processed RNA-seq data; KW, XL, SLS, YC analyzed results; SLS, AC, YC revised manuscript; YC wrote the manuscript, supervised the project and secured funding.

---

**Figure S1. iRIL-seq library analysis.**

A) Hfq iRIL-seq Data are enriched with sRNAs and mRNAs. Single (Single), chimeric (Chimera) and statistically significant chimeric (S-chimera) sequenced fragments from Hfq-Flag and IgG libraries were classified into eight major categories: 5’ UTR, CDS, 3’ UTR, sRNA, tRNA, rRNA, hkRNA (house-keeping non-coding RNAs include SRP RNA and RNase P RNA), IGR (intergenic region). Total counts are denoted in parentheses.

B) Heat map showing that iRIL-seq replicates were highly reproducible at the single fragment counts and the chimeric fragment counts level.

**Figure S2. Consensus motifs identified in target mRNAs by iRIL-seq**

Motif found in target sequence set of each sRNA that had at least ten targets, including RprA, CpxQ, FnrS, SdsR, CyaR, GcvB, SdhX, ChiX, ZbiJ, MicL, DapZ and FadZ.

**Figure S3. The overexpression of ArcZ and its effects on HMV.**

A) Analysis of HMV in the absence of IPTG.

B) The expression of ArcZ was induced by IPTG. Leaky expression of ArcZ was visible in the absence of IPTG.

C) ArcZ overexpression reduced the amounts of capsule polysaccharides in HvKP. Capsule was extracted from bacterial cultures and analyzed by 4-12% SDS-PAGE and Alcine blue staining. Capsule extracted from the acapsular i-*wza* mutant was used as a negative control.

**Figure S4. Putative LacI-like repressors in the genome of ATCC 43816.**

A) BlastP results showing the homologs LacI repressors in the *K. pneumoniae* ATCC 43816 genome, using *E. coli* LacI as query.

B) Multiple alignment of LacI-like protein sequences. The putative DNA-binding domain is boxed.

**Figure S5. ArcZ represses HMV independent of *arcB*.**

A) Confirmation of the *arcZ* deletion strain and the ArcZ chromosomal complementation strain. The expression of mature ArcZ sRNA was detected by northern blot using probe YCO-1745. 5S rRNA serves as the loading control.

B) HMV was fully repressed by ArcZ in the D*arcB* mutant. HMV was determined for each strain in the presence of IPTG and hygromycin overnight. Relative HMV levels were shown with WT strain set as 1. pJV300 was used as control vector and pZE12-ArcZ was used to overexpress ArcZ in the presence of IPTG. The i-*wza* mutant was used as the acapsular negative control.

**Figure S6. Analysis of the ArcZ-interacting mRNAs in ATCC 43816**

A) Gene ontology analysis showing that ArcZ targets were enriched in carbohydrate metabolism and polysaccharide metabolism.

B) Browser image showing the ArcZ chimeras mapped in the capsule gene cluster. For each ArcZ-targeting capsule gene, the count of chimeric fragments is denoted in parentheses.

**Figure S7. ArcZ downregulates the expression of capsule genes.**

The expression of capsule genes and *rmpACD* regulators was determined by RNA sequencing. The relative mRNA levels were shown in relative to one of WT+pJV300 samples based on CPM (counts per million reads). The experiment was performed in biological triplicates.

**Figure S8. ArcZ overexpression inhibits target-GFP reporters.**

A) The MlaA-sfGFP reporter was repressed by WT ArcZ, but not by ArcZ-M2 and ArcZ-M5 mutants. GFP was detected using Western blotting with an anti-GFP antibody; GroEL serves as the loading control.

B) The Fbp-sfGFP reporter was repressed by WT ArcZ, but not by ArcZ-M2 and ArcZ-M5 mutants.

C) The control GFP reporter was not affected by ArcZ overexpression.

D) Northern blotting analysis confirmed the overexpression of ArcZ variants. Mature ArcZ was detected using probe YCO-1745. 5S rRNA serves as the loading control.

**Figure S9. ArcZ may be a repressor of *Klebsiella* virulence.**

Six mice per group were infected WT, Δ*arcZ* mutant and an *arcZ* complementation strain containing pZE12-*P_own_-arcZ*), respectively. Mice were scarified 48 h post-infection to determine the bacterial burden in organs (CFU/ml). *, p < 0.05; **, p < 0.01; ns: not significant, Mann-Whitney test.

**Figure S10. ArcZ overexpression inhibits *Klebsiella* virulence in mice.**

A) ArcZ overexpression significantly reduced bacterial burden in the lungs of infected mice. Mice were infected with 10^3^ CFU of the indicated strains, and scarified at 48 hpi. *, p < 0.05; **, p < 0.01; ns: not significant, Mann-Whitney test. The 16S rRNA P1 promoter was used to overexpress ArcZ from plasmid in the WT strain (WT+pZE12-*P_16S_-arcZ*); the *i-wza* acapsular mutant was used as a negative control.

B) The Δ*mlaA* mutant was outcompeted by WT HvKP in mice. Mice were infected with a 1:1 mixture of WT and Δ*mlaA* strains in a total inoculum of 10^3^ CFU via intranasal instillation. After loss of 10% of body weight, mice were sacrificed and the competitive index was determined by measurement of viable counts in mice lungs. *, p < 0.05; **, p < 0.01; Mann-Whitney test. The *in vitro* competitive index was determined after overnight growth in LB and CFU numeration.

**Figure S11. ArcZ is activated catabolite repression and CRP**

A) Northern blot analysis showing ArcZ levels reduce upon addition of 0.5% glucose in LB. The CyaR sRNA was probed as a positive control, as an established CRP-dependent sRNA. 5S rRNA served as a loading control. The pre-ArcZ transcript was detected using the probe YCO-1838, and mature ArcZ was detected using probe YCO-1745. 5S rRNA is the loading control.

B) Northern blot analysis suggests that ArcZ was not regulated by *arcA/arcB* in *K. pneumoniae*. Wildtype cells and isogenic derivatives deleted for *arcA* or *arcB* were grown overnight in presence or absence of oxygen. The pre-ArcZ transcript was detected using the probe YCO-1838, and mature ArcZ was detected using probe YCO-1745.

**Figure S12. ArcZ acts downstream of CRP to repress HMV.**

A) Both ArcZ and CRP inhibit HMV. After overnight growth in LB, the HMV was determined in the indicated mutant strains, with the *i-wza* mutant as negative control. N=3 biological replicates, bars indicate mean ± SD. *, p < 0.05; **, p < 0.01; Student’s t-test.

B) ArcZ moderately reduced HMV in the absence of CRP. After overnight growth in LB+IPTG, the HMV was determined in the indicated strains with pJV300 or pZE12-ArcZ plasmids. The *i-wza* mutant was included as negative control. N=3 biological replicates, bars indicate mean ± SD. *, p < 0.05; **, p < 0.01; Student’s t-test.

**Table S1** Illumina sequencing reads statistics

**Table S2** *Klebsiella* sRNA annotations and enrichment in Hfq-coIP

**Table S3** Enrichment of *Klebsiella* mRNAs in Hfq-coIP

**Table S4** Complete list of significant RNA-RNA interactions in *K. pneumoniae*

**Table S5** The list of HMV-related genes in *K. pneumoniae*

**Table S6** Interactions with HMV-related genes in *K. pneumoniae*

**Table S7** Clinical isolates used in this study

**Table S8** Bacterial strains used in this study

**Table S9** Plasmids used in this study

**Table S10** Oligonucleotides used in this study

## Notes

### Competing Interest Statement

The authors have declared no competing interest.

